# IT IS NOT A SMALL WORLD FOR PSYCHIATRIC PATIENTS: Small-world of psychiatric patients

**DOI:** 10.1101/2024.03.25.586529

**Authors:** Ata Akın, Emre Yorgancıgil, Ozan Cem Öztürk, Bernis Sütçübaşı, Ceyhun Kırımlı, Elçim Kırımlı, Seda Nilgün Dumlu, Gülnaz Yükselen, S. Burcu Erdoğan

## Abstract

Individuals suffering from Obsessive Compulsive Disorder (OCD) and Schizophrenia (SCZ) frequently exhibit symptoms of cognitive disassociations, which are linked to poor functional integration among brain regions. The loss of integration can be assessed using graph metrics computed from functional connectivity matrices (FCMs) derived from neuroimaging data. A healthy brain with an effective connectivity pattern exhibits small-world features with high clustering coefficients and shorter path lengths in contrast to random networks. We analyzed neuroimaging data from 60 subjects (13healthy controls, 21 OCD and 26 SCZ) using functional near-infrared spectroscopy (fNIRS) during a color word matching Stroop Task and computed FCMs. Small-world features were evaluated using the Global Efficiency (*GE*), Clustering Coefficient (*CC*), Modularity (*Q*), and small-world parameter (*σ*). The proposed pipeline in this study for fNIRS data processing demonstrates that patients with OCD and SCZ exhibit small-world features resembling random networks, as indicated by higher *GE* and lower *CC* values compared to healthy controls, implying a higher operational cost for these patients.

**AUTHOR SUMMARY:** Individuals suffering from Obsessive Compulsive Disorder (OCD) and Schizophrenia (SCZ) frequently exhibit symptoms of cognitive disassociations, which are linked to poor functional integration among brain regions. The loss of integration can be assessed using graph metrics computed from functional connectivity matrices (FCMs) derived from neuroimaging data. A healthy brain with an effective connectivity pattern exhibits small-world features with high clustering coefficients and shorter path lengths in contrast to random networks. We analyzed neuroimaging data from 60 subjects (13healthy controls, 21 OCD and 26 SCZ) using functional near-infrared spectroscopy (fNIRS) during a color word matching Stroop Task and computed FCMs. Small-world features were evaluated using the Global Efficiency (*GE*), Clustering Coefficient (*CC*), Modularity (*Q*), and small-world parameter (*σ*). The proposed pipeline in this study for fNIRS data processing demonstrates that patients with OCD and SCZ exhibit small-world features resembling random networks, as indicated by higher *GE* and lower *CC* values compared to healthy controls, implying a higher operational cost for these patients.

## INTRODUCTION

The human brain is a complex network made up of millions of neurons with a small world topology, which is widely used to model the network behavior of various natural interactions involving social networks and brain networks (Achard & Bullmore, 2007; Bassett & Bullmore, 2006; Bullmore & Sporns, 2009; Rubinov & Sporns, 2010).

The understanding of brain network function through its correspondence to small-world topology is predicated on the assumption that information processing and transfer between neuronal structures have an optimal dynamic balance of local-level specialization and global-level integration. The small-world topology is defined by dense local clustering of neuronal connections between neighboring nodes and a short average path length between all possible pairs of nodes (Bassett & Bullmore, 2006). The functional and structural organization of the human cerebral cortex have been frequently modeled by a small-world topology because the properties of a small-world configuration optimize both local information processing efficiency and global information integration, where efficiency is defined as low energy consumption and wiring costs with a high transmission rate.

Furthermore, the application of computing small-world topology features to the interpretation of structural and functional neuroimaging data obtained at various spatial and temporal scales has demonstrated that any disruption in the structural organization or functionality of this network could be an indicator or a result of a brain disease or disorder. Indeed, several neuropsychiatric disorders have been linked to disruptions and changes in the small-world properties of cerebral cortex networks. Among these disorders, changes in functional and structural connectivity have been extensively studied for obsessive-compulsive disorder (OCD) and schizophrenia (SCZ). Resting-state functional magnetic resonance imaging (fMRI) studies have revealed that SCZ patients have lower small-world functional connectivity and weaker local clustering (Andreasen et al., 1999). Deficit and nondeficit SCZ patients have lower local efficiency metrics than healthy controls, indicating a decrease in local specialization (M. Yu et al., 2017).

The efficiency of a network’s organization can be measured using graph theoretical metrics. Previous studies on brain functional connectivity have benefited significantly from graph theoretical metrics in generating such features (Bullmore & Sporns, 2009; Deuker et al., 2009; Klados et al., 2013; Niu et al., 2013). Among many metrics, Global Efficiency (*GE*) has been widely and consistently used in brain connectivity research as a metric to quantify the information-sharing efficiency of functional connectivity matrices (FCMs)(Adolphs, 2003a, 2003b, 2009; Akin, 2017; Deuker et al., 2009; Klados et al., 2013; Palva, Monto, & Palva, 2010). Small-world phenomena imply both high local and global efficiency, which manifests itself as high functional specialization measured by a metric named the clustering coefficient and cost-efficient functional integration measured by a shorter mean path length (Bassett & Bullmore, 2006; Bullmore & Sporns, 2009).

Over the last two decades, numerous studies have focused on how these graph theoretical metrics can be computed from neuroimaging data and how well they represent brain states, particularly the affected brain. Many studies investigated the accuracy of such metrics as biomarkers extracted from neuroimaging data and discovered conflicting results for SCZ patients (see (Gao et al., 2023) for a recent meta-analysis). As a less studied disease using graph theoretical approaches, OCD patients showed more consistent results with *GE* than healthy controls (Akin, 2021; Li, Li, Cao, et al., 2022; Li, Li, Jiang, et al., 2022; Thorsen et al., 2021).

Overall, studies found an impairment of small-world organization of the cerebral functional network in psychiatric patients than healthy controls. This study investigates the small-world properties of prefrontal cortical functional networks in psychiatric patients to challenge previous research findings. We investigate the use of fNIRS, a rapid, non-invasive, mobile brain imaging technology that is better suited to cognitive studies, in elucidating the small-world properties of prefrontal cortical functional networks.

## METHODS AND MATERIALS

### Subjects and Experimental Procedure

The study included sixty subjects from three different groups. The three categories were as follows: 1) 13 healthy control subjects (6 female, average age 26), 2) 26 patients with obsessive-compulsive disorder (11 female, average age of 29), and 3) 21 patients with SCZ (10 female, average age of 28). Diagnoses were made based on comprehensive interviews conducted by an experienced psychiatrist using standardized diagnostic criteria (DSM-IV-TR). The protocol was approved by the Ethics Committee of Pamukkale University in 2008. Our group has previously published portions of this data using analysis methods other than those used in the current study (Akgul, Akin, & Sankur, 2006; Ata Akin et al., 2006; Ciftçi, Sankur, Kahya, & Akin, 2008; Dadgostar, Setarehdan, Shahzadi, & Akin, 2016; Dalmis & Akin, 2015; Einalou, Maghooli, Setarehdan, & Akin, 2015, 2016, 2017; Erdoğan & Yükselen, 2022; Koray Ciftci, Bulent Sankur, Yasemin P Kahya, & Ata Akin, 2008; Sergul Aydore, M. Kivanc Mihcak, Koray Ciftci, & Ata Akin, 2010). All subjects gave written and signed consent to participate in the study before the experiment. The subjects were instructed to perform the task while sitting in front of a computer in a dimly lit, isolated room.

Subjects were asked to respond to stimuli for a computerized color word-matching Stroop task, which has been thoroughly described in previous research (Koray Ciftci et al., 2008; Zysset, Müller, Lohmann, & von Cramon, 2001). The subjects were instructed to respond with left mouse clicks with their index finger for “YES” and right mouse clicks with their middle finger for “NO” with their right hand, depending on whether the stimulus was correct or incorrect. During the “Neutral” trials, the top row showed letters (“XXXX”) in different colors (red, green, or blue), while the bottom row showed color words (“RED,” “GREEN,” and “BLUE”) printed in black. In the “Congruent” trials, the top row featured color words printed in matching colors. Conversely, in “Incongruent” trials, color words were printed in opposing colors to induce interference. Participants were asked to determine whether the color name in the top row matched the color word in the bottom row under all conditions. Half of the trials in each condition showed colors that were consistent between the top and bottom rows. The stimuli were presented in nine randomly assigned blocks, with each block containing five consecutive stimuli of the same type. The blocks were interspersed with 20-second rest intervals. Each stimulus within a block was displayed on the screen for 2.5 s, with an inter-stimulus interval of 4 s. The subjects had to respond with left or right mouse clicks based on whether the stimulus was correct or incorrect. The entire Stroop task began with 30 s of resting fNIRS recording and concluded with another 30 s of rest.

### Cognitive Quotient

The Stroop task yields two types of behavioral outcomes: Reaction (Response) Times 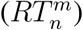 in seconds and Accuracy 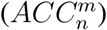 as a percentage value, concerning task type *n* = *N, C, I*, for each subject *m*. RTs are calculated by averaging the response times of correctly answered stimuli. Accuracy was calculated as the ratio of total correct answers to total questions. Combining these two parameters yields the Cognitive Quotient 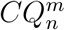, also known as cognitive efficiency), which can be calculated by 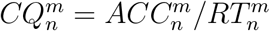. *CQ* can also be viewed as a measure of cognitive behavior that includes both reaction time and error rates.

### fNIRS Equipment

A custom-built fNIRS system (NIROXCOPE 301) was used to collect cortical hemodynamic data at a sampling frequency of 1.77 Hz from 16 brain regions using a flexible probe placed over the forehead. The NIRS system includes a flexible probe placed on the subjects? foreheads, a data acquisition unit, and a data collection computer. The prefrontal cortex probe has four dual-wavelength light-emitting diodes (LED, *λ*_1_ = 730 nm and *λ*_2_ = 850 nm) and 10 photodetectors, forming 16 equidistant light source and detector pairs at a distance of 2.5 cm (Figure 1). A channel is defined as a pair of 1 LED and 1 photodetector separated by 2.5 cm. Several tests of the validity of this instrument and probe design for probing brain tissue (Erdoĝan, Yücel, & Akin, 2014), as well as its ability to detect cognition-related brain signals, were provided in our previous studies (Akin, 2017; Ciftçi et al., 2008; Dadgostar et al., 2016; Dalmis & Akin, 2015; Einalou, Maghooli, Setarehdan, & Akin, 2014; Sergul Aydore et al., 2010).

**Figure 1.**
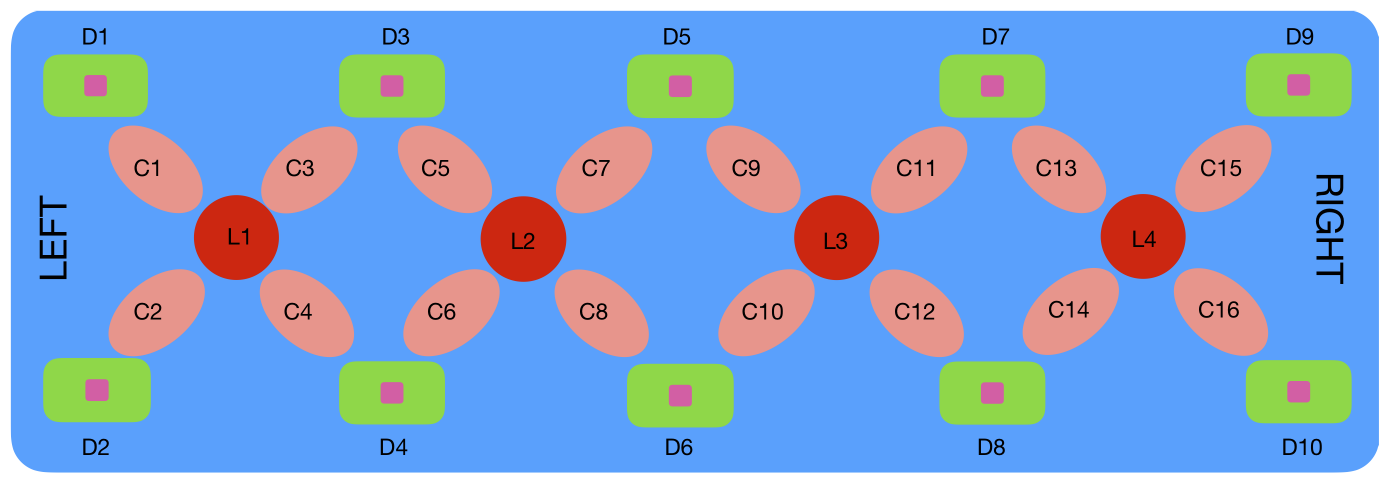
Rectangular probe geometry of the fNIRS NIROXCOPE 301. *L*_*i*_ are the LEDs, *D*_*i*_ the detectors, and *C*_*i*_ the channels (Ata Akin et al., 2006; Aydöre et al., 2010).

### Analysis of the fNIRS Data

An overview of the analysis of fNIRS [HbO] data consisted of 7 major steps, which will be detailed in the following sections. Briefly, the analysis steps included: 1) Calculating partial correlation-based FCMs for both rest and task-stimulus intervals, 2) Calculating the average of FCMs across all subjects and stimulus types to find one common background FCM and subtracting this background from each task-based FCM (Background Subtracted: BS), 3) performing PCA on the BS FCMs and reconstructing noise-eliminated FCMs, 4) Applying a thresholding procedure to BSPCA FCMs, 5) Computing the *GE* value from each combination of PCA set and threshold, 6) Determine the PCA component set and threshold with the highest statistically significant difference among the three subject groups based on the *GE* parameter, and 7) Calculate the *CC, Q* and *σ* parameters of the small world phenomena using the selected PCA components and thresholds from step 6. Figure 2 shows the flow of the signal processing pipeline. The sections that follow explain the chronological set of steps in this pipeline *Preparing FCMs* Our previous research confirmed the benefits of using a partial correlation-based analysis to obtain FCMs (Akin, 2017, 2021; Einalou et al., 2017). Similar to previous studies, 16 channels of fNIRS [HbO] time series data were first high-pass filtered with a Butterworth filter of order 8 at a cutoff frequency of 0.09 Hz. The subject-based average of these high-pass filtered data was then used as the regressor in the partial correlation analysis, as described in (Akin, 2017, 2021). In summary, the partial correlation coefficient between each pair of channels (*i, j* = 1 … 16) can be calculated using the raw (unfiltered) [HbO] data (*k*) from two channels with the mean regressor as follows:

**Figure 2.**
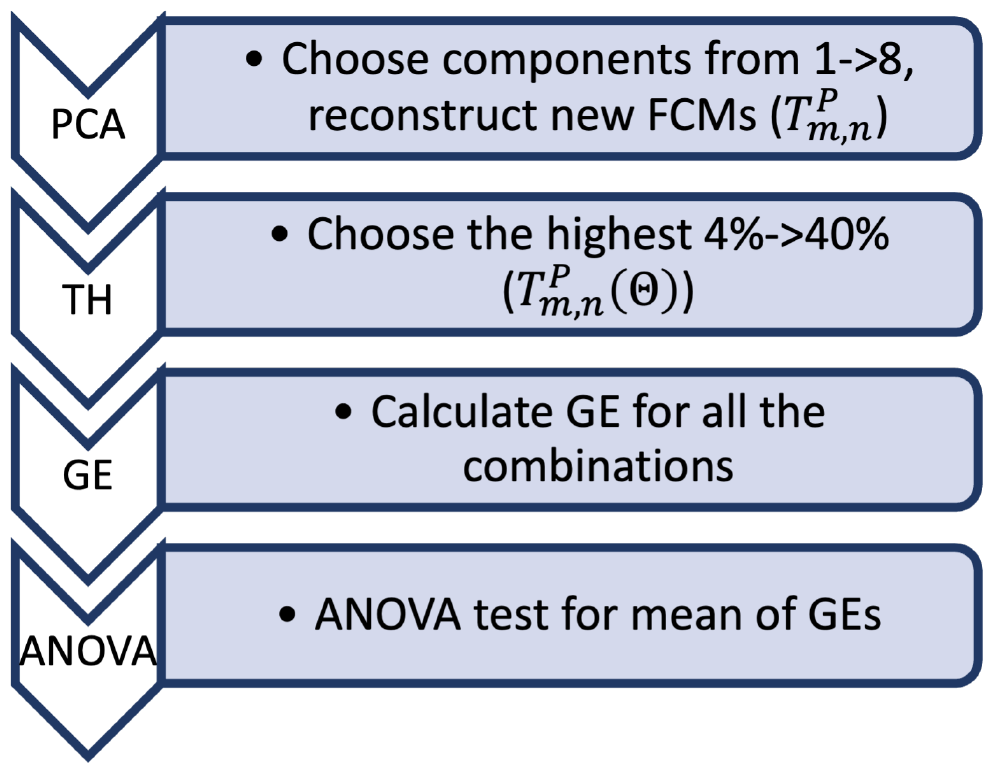
Algorithm of the proposed work: PC-FCMs: Partial correlation-based FCMs: Partial correlation based FCMs (Eq. 1), BS: Back Subtraction, PCA: PCA to obtain Task Positives FCM (**T**_*P*_, Eq. 6), TH: Thresholding and computing adjacency matrix (𝕋_*P*_ (Θ)), ANOVA: Calculate 3-Class statistical significance (*p*-value)

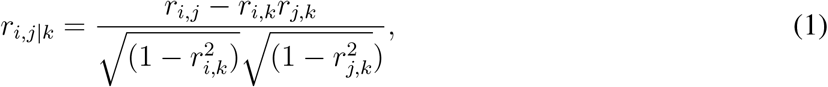

Raw [HbO] data were concatenated according to task type and rest data. Hence, from one channel of fNIRS data, four concatenated data sets were generated (1 for resting-state and 3 for task-based). In total, 16X4 data traces were created for each subject, which were then used to generate 4 FCMs. Because there are 16 channels, the FCM is converted to a 16x16 symmetric matrix with diagonals nulled (*r*_*i,i*|*k*_ = 0) to exclude them from further *GE* analysis. In this way, 4 FCMs are generated for each subject using the four sets of concatenated signals.

*Background Network Subtraction* In addition to the previous studies, the question of whether FCMs have background network connectivity that obscures the task-related network topology that emerges during cognitive tasks was raised. We decided to test this assumption by asking a simple but fundamental question: How much do resting-state FCMs and task-based FCMs resemble each other?

To answer this question, the average of the remaining FCMs 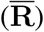 and task-based FCMs 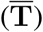 was calculated across all 60 subjects. The hypothesis to be tested was:

**Assumption 1** *A background connectivity network is active even during cognition, which obscures the cognitive network structure*.

Therefore, the following average of the rest- and task-based FCMs over 60 subjects were calculated to test for this assumption:

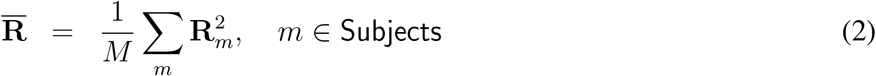

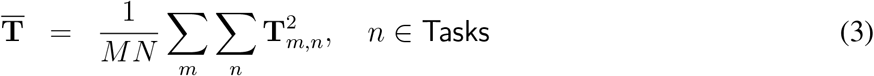

where *m* = 1 … *M* (*M* = 60 number of subjects) and *n* = 1 … *N* (*N* = 3). Note that the averages were calculated using squared FCMs. The FCMs are squared for the remainder of the analysis. Averaged FCMs for all subjects during rest and task episodes are shown in Figures 3(a) and 3(b). Here, 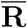 represents the average resting state of squared FCMs calculated across all subjects, while 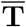 represents the averaged task-based squared FCMs calculated across all subjects.

**Figure 3.**
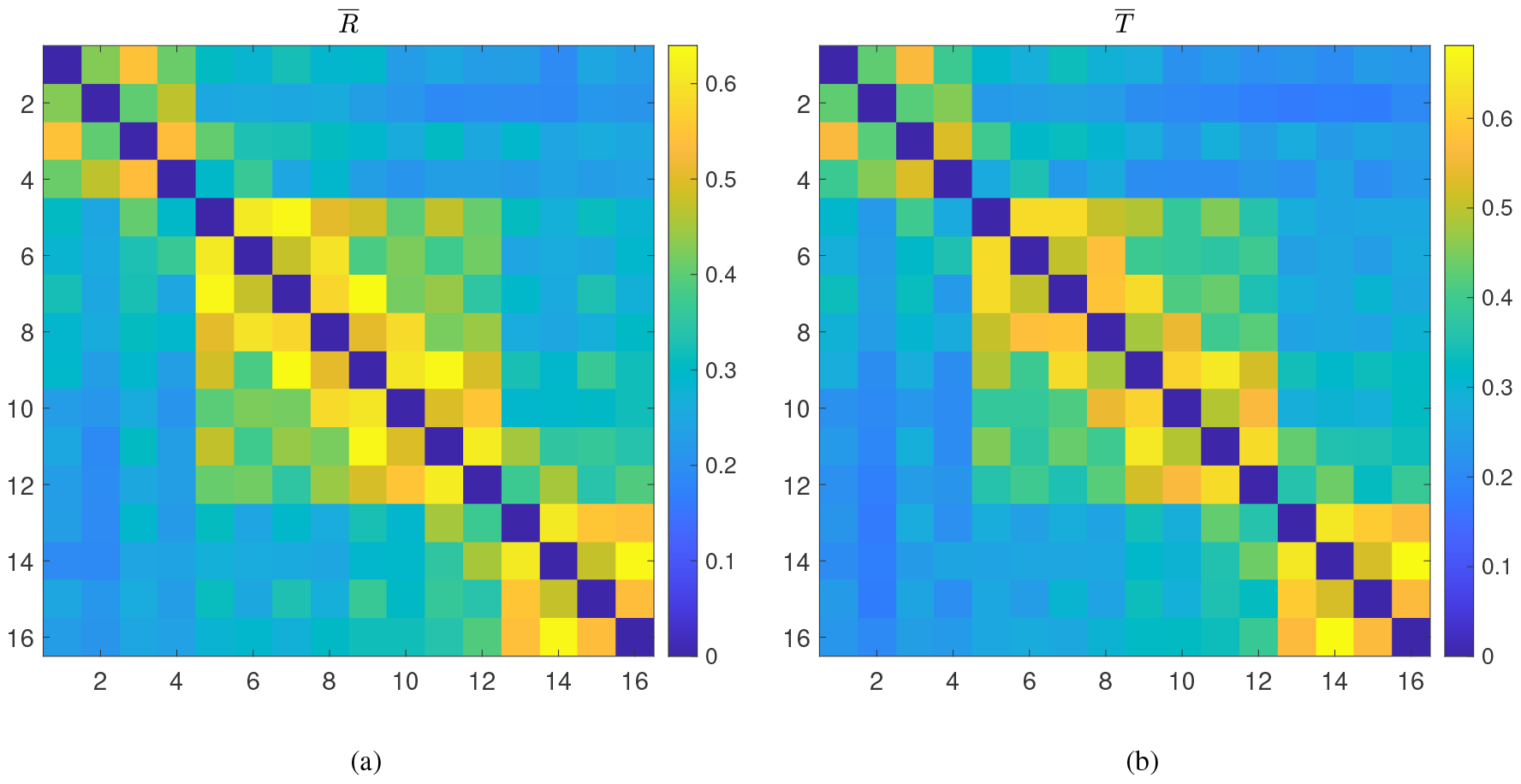
a) 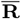, b) 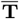

A significant correlation value computed using 2-dimensional correlation analysis (*r* = 0.9941, *p <<* 10^*−*10^) was found between these two matrices. Therefore, it is now clear that even partial correlation analysis was insufficient to reduce the dominance of physiological systemic activity observed in the background network over networks active during cognition. The presence of a dominant background network in the task FCMs necessitates the removal of this background network, which could not have been eliminated using partial correlation analysis alone. In our previous paper (Akin, 2021), we did not perform background subtraction because we assumed that the PCA-based approach would already eliminate all background task-unrelated connectivity topologies. These matrices will now be treated as images, and the following processing pipeline will be based on image-processing approaches developed for target and motion detection applications (Oliver, Rosario, & Pentland, 2000; Piccardi, 2004).

Previous research on removing static background images from images containing moving or non-background targets proposed subtracting this background to improve target detection accuracy (Elgammal, Harwood, & Davis, 2000; Horprasert, Harwood, & Davis, 1999). The following processing steps are performed to remove the static background FCM from the task-based FCM:

1. Determine the common background from the average of the *rest* FCMs 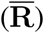 from Equation 2 and the task-based FCMs 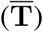 from Equation 3, and call it the background connectivity FCM 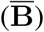:

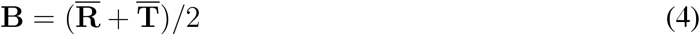
2. Subtract the common background from the task-based FCM (**T**)

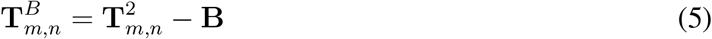

where *m* denotes the subject, and *n* represents the stimulus type. The assumption behind averaging across all subjects is that there is a common baseline resting-state network in all human brains, that remains active during any cognitive paradigm and contains an intrinsic functional connectivity pattern (unrelated to sensory or motor stimuli) that is responsible for maintaining bodily functions (Fox et al., 2005; Raichle et al., 2001; Ralchle & Snyder, 2007). Cognition or task-based FCM, denoted as **T**_*m,n*_ cn can be assumed to be superimposed on this previously existing so-called “*evolutionary network*” (**B**) (Hofman, 2012, 2014; Yeo et al., 2011). Therefore, while the brain is doing what it always does (maintaining all bodily functions, as evolution dictates), it should also focus on a specific cognitive task. Once background subtraction is completed, task-based FCMs with small-world properties can be easily extracted. Further elimination of background noise can be done using PCA, which has been proposed in image processing applications (Nandi et al., 2015; Rodarmel & Shan, 2002). **Assumption 2** *Cognition necessitates the activation of an* independent *functional neural network called the task-positive Network (***T***P)*
3. Perform PCA on the 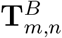
4. Reconstruct a new Task Positive Network **T**^*P*^ with the *strongest* components of the PCA decomposition (Xia, Wen, Eberl, Fulham, & Feng, 2008)

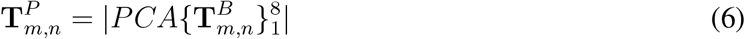

This pipeline generated 3 new task-based reconstructed FCMs for each subject. Assuming that these FCMs are now free of background “blurring” effects, we can proceed with the generation of features from these matrices.

A thresholding algorithm was applied to task-positive FCMS, in which only a subset of the highest correlation values within a single matrix were retained to ensure that the same number of edges appeared across all FCMs. This threshold value eventually determines how many nonzero elements are present in the binaries matrix, also known as the adjacency matrix.

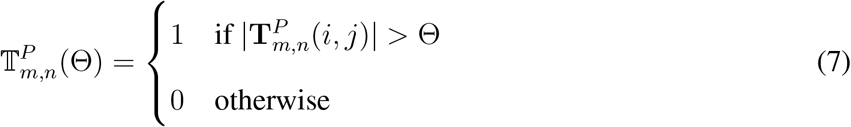

Eventually, we are left with metrics for a specific combination of PCA components (*i*.*e*. the first 3) and a threshold value. The threshold value determines how sparse the matrix is. Sparsity is calculated by dividing the number of nonzero elements in a matrix by the total number of elements in a matrix. The sparsity (also known as the *Cost*) values were chosen between 4% and 40%.

### Small-Worldness

Small-world phenomena in brain networks can be studied by comparing graph theoretical metrics obtained from individual FCMs concerning different threshold values. Typically, a comparison is performed between several of the graph theoretical metrics, such as the *GE*, Clustering Coefficient (*CC*), characteristic path length (*L*), and graph theoretical modularity (*Q*) values, with those obtained from a normally distributed random matrix (network) and those from a lattice type of network for varying levels of sparsity or cost, as described in (Achard & Bullmore, 2007; Alexander-Bloch et al., 2010).

Computation of the graph theoretical metrics is explained in detail in other studies (Bullmore & Sporns, 2009; He & Evans, 2010; Skidmore et al., 2011). The small-world parameter (*σ*) can be calculated from the other metrics using the following formula (Bassett & Bullmore, 2017):

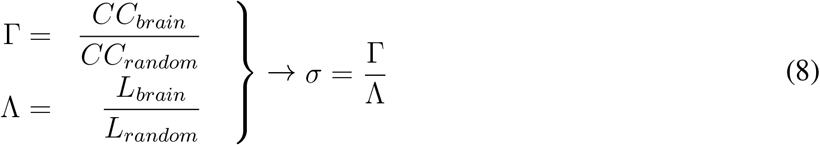

These equations were used to calculate the individual *σ*^*C*^ for controls, *σ*^*O*^ for OCD patients, and *σ*^*S*^ for SCZ patients. Therefore, the primary goal of this paper is to test the following hypothesis:

**Hypothesis 1** *The small-world phenomenon is disrupted in OCD and SCZ patients due to changes in the brain’s network of functional connectivity*.

A comparison was made between the small world parameters of *GE, CC, Q*, and *σ* values of the subjects to test the above hypothesis.

## RESULTS

### Behavioral Results

Figure 4(a) shows the average *CQ* values for subjects and stimuli. A two-way mixed ANOVA of mean *CQ* values for each group and stimulus type ((3)*GROUPS* × (3)*STIMULUS*) revealed significant values for group comparison (*p <* 10^*−*7^), and stimulus type (*p <* 10^*−*7^) but no interaction for *GROUPS * STIMULUS* (*p* = 0.97). There is a decreasing trend of the mean *CQ* as the task becomes more difficult (from N to I) for all groups. Similarly, the mean *CQ* value 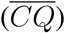 plotted as the last bar for each group across three stimuli types (shown in Figure 4(b)) resulted with 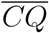 as 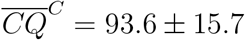 for controls, 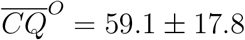 for OCD patients, and 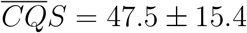 for SCZ patients with a significant difference (*p* = 1.26 exp −10). Because there is no significant interaction for this term, the 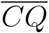 (over three stimulus types) will be used for the remainder of the analysis.

**Figure 4.**
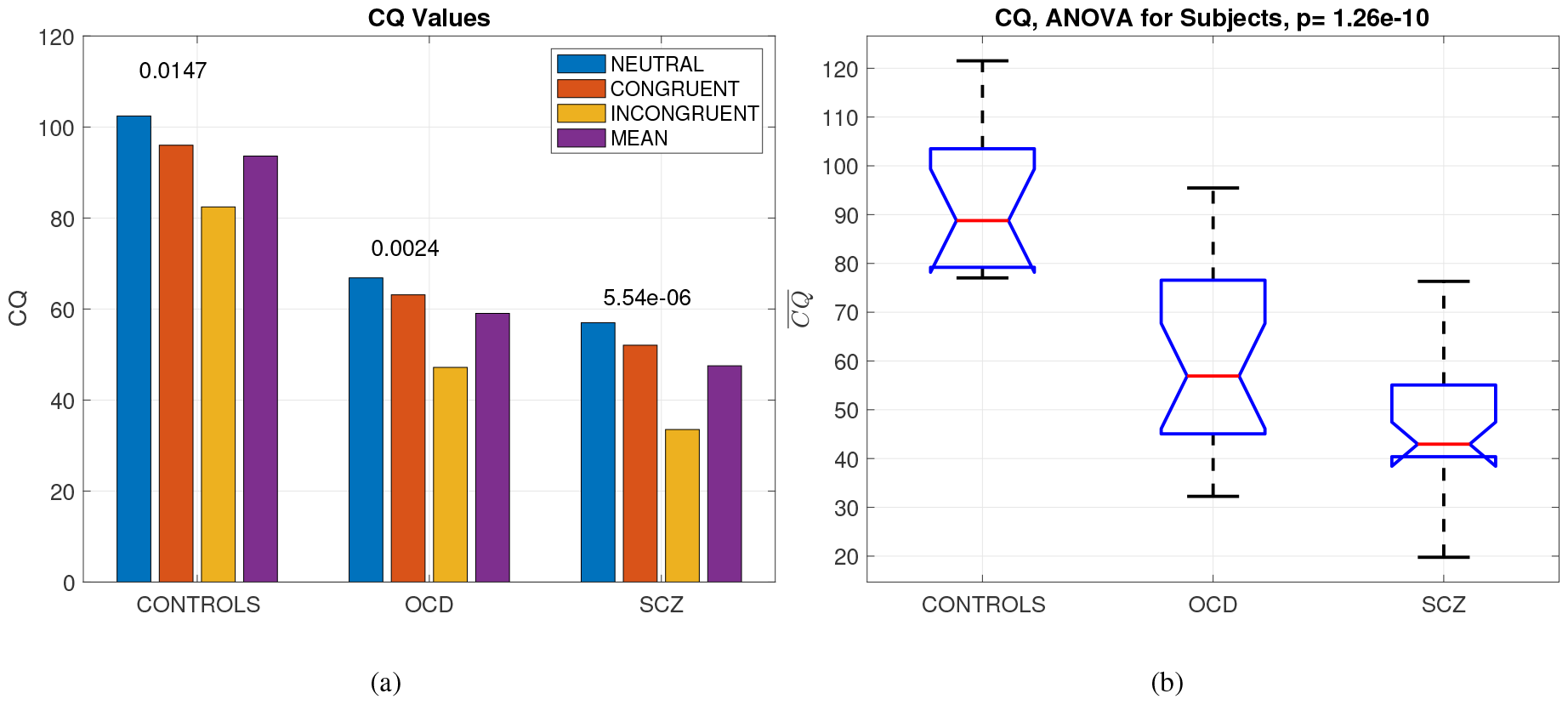
(a) *CQ* values for groups and task types, with within-group p values shown on top of the bars and the *p* value between subjects for mean CQ values over three tasks in the title. (b) ANOVA for the 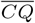values for groups observed as purple bars in (a).

### Not such a small-world for psychiatric patients

There are several ways to determine whether a network (or an FCM) exhibits the small-world phenomenon. One approach is to compare subjects’ graph-theoretic metrics to values from random and lattice-type matrices (Achard & Bullmore, 2007; Bassett & Bullmore, 2017; Bassett et al., 2009). 60 random FCMs with normal distribution were generated, and their graph-theoretic metrics were calculated using the same thresholding method described previously. Similarly, 60 lattice-type FCMs were created using the code `(makelatticeCIJ(16, TH))` from the Brain Connectivity Toolbox(Rubinov & Sporns, 2010).

Figure 5 shows the violin charts for the *GE, CC, Q*, and *σ* values at different thresholds and compares them to random and lattice networks. The threshold interval was selected based on the statistics with the highest significance (*p <* 0.05) for selecting the PCA set (ℙℂ) and threshold values (**Θ**) of 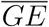

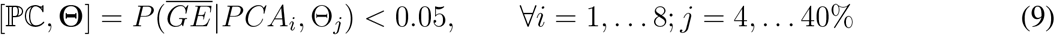

For this dataset, the optimal PCA components were ℙℂ = [1 → 6] and **Θ** = [10.5 → 19.5]%. As predicted, in Figure 5(a), human *GE* values present a small-world phenomenon, because they fall between the two extreme cases of network topologies, namely the lattice and random networks. As observed, the values of people with OCD and SCZ have values that are closer to random networks, with SCZ being closer to randomness. This pattern also appears in the other three metrics. Figure 5(b) displays the distribution of *CC*, 5(c) the modularity *Q*, and finally 5(d) the change in the small-worldness index, *σ*. All of the graph’s theoretical metrics indicate a statistically significant shift toward randomness in people with OCD and SCZ.

**Figure 5.**
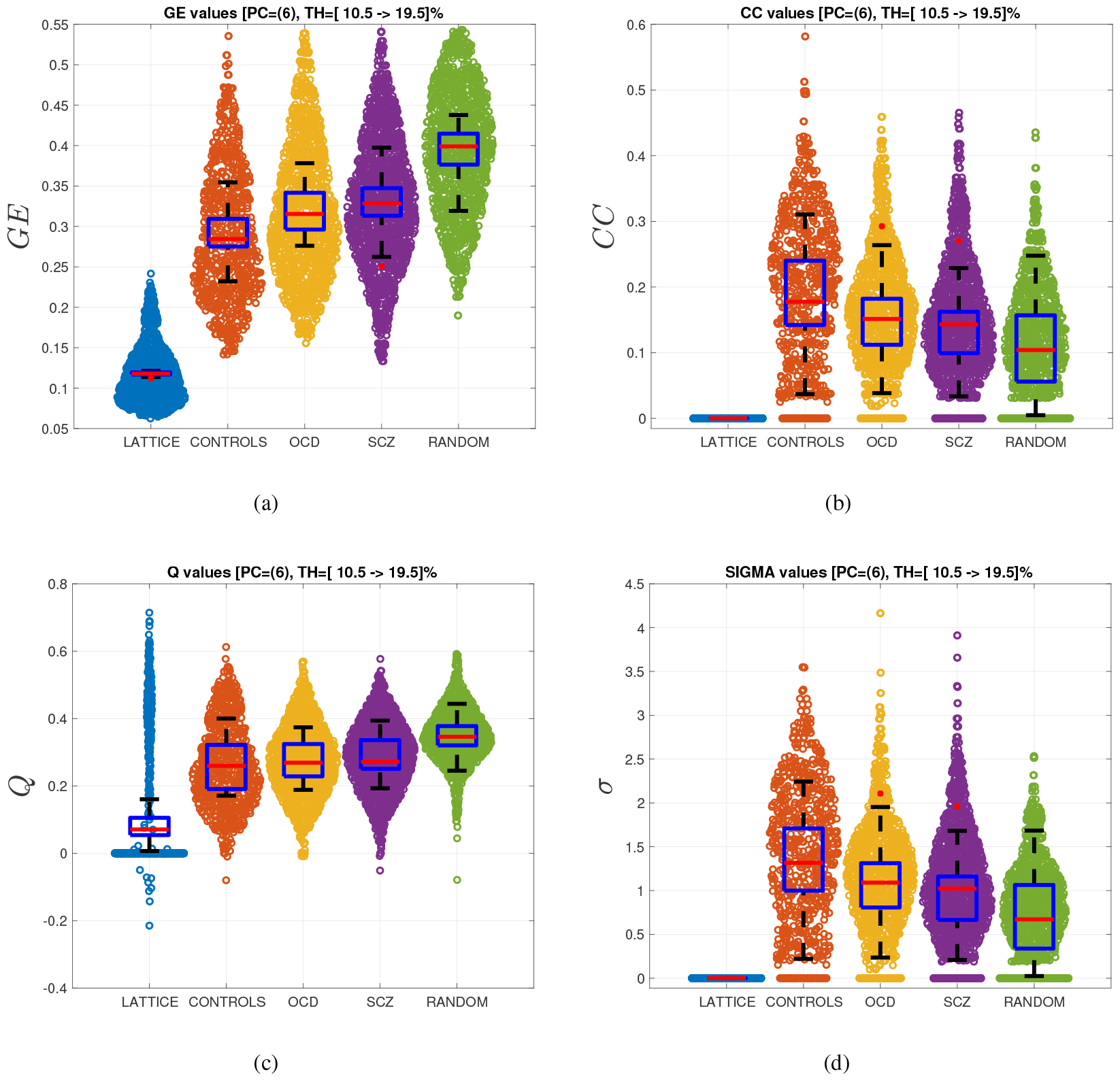
Violin charts for (a) 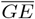, (b)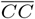, (c) 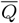,and (d) 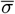 The boxplots are derived for the mean values over the thresholds.

The boxplots in the center of the violin charts are calculated using the average of the individual charts. We present these boxplots to demonstrate how the medians and standard deviations differ between groups.

### Connectivity Patterns

Connectivity maps were created by averaging all the task-based FCMs after BS&PCA processing for both healthy controls and affected subjects. When the averaged FCMs were computed, the threshold that gave the highest statistical significance for the 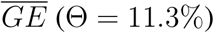 was applied to these two matrices to identify the remaining connections. Figure 6(a) illustrates the final connectivity map (*i*.*e*. adjacency matrix) for the controls with an overall 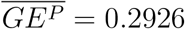, while Figure 6(c) depicts the affected subjects, with 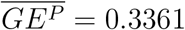 . Healthy controls and affected subjects use different nodes when performing the Stroop task. Healthy controls used the middle to right prefrontal cortex more frequently, whereas affected subjects used the opposite pattern.

**Figure 6.**
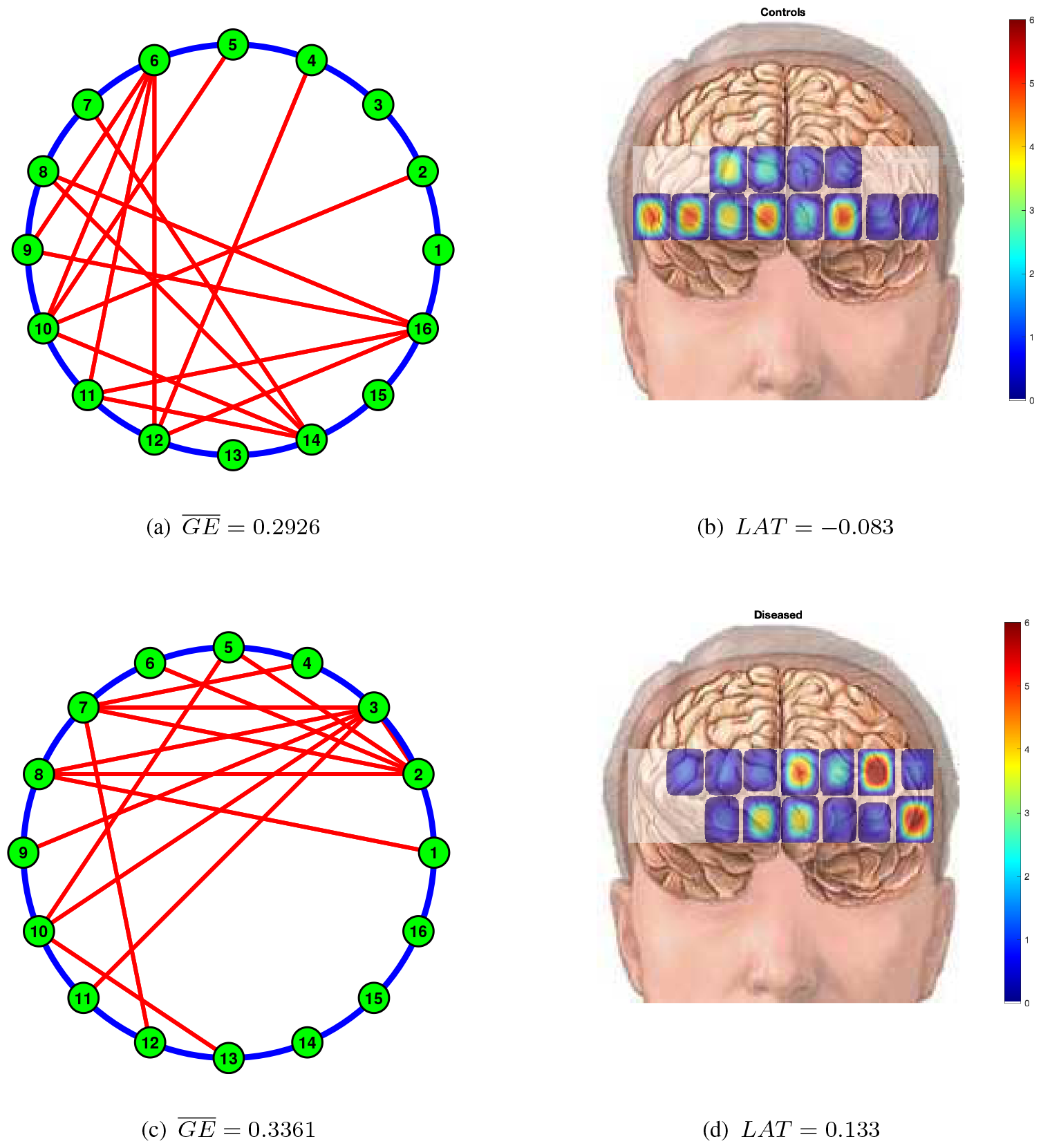
Connectivity maps (a) for healthy controls, and (c) affected subjects and the connectivity strength maps for (b) healthy controls and (d) affected subjects.

Another way to display these connectivity maps is by calculating the strength of connections for each node (*i*.*e*. the sum of connections from a node using the adjacency matrices). A laterality index can be defined to evaluate the hemispheric dominance of the connections as follows:

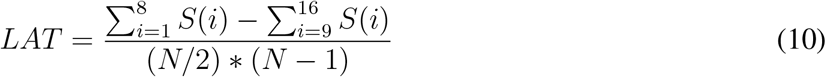

where *S*(*i*) is the total number of connections from the *i*^*th*^ node, *N* = 16. The *LAT <* 0 indicates right lateralization, and vice versa. The *LAT* values quantify the brain’s hemispheric dominance.

### Neuroscientific Results

Neuroscientific research seeks to understand the relationship between neurobiology and behavior. Typically, this relationship is assumed to be casual. When biology changes, behavior should follow suit.

This trend is typically expressed as a correlation between behavioral outcomes and biological findings, as illustrated in Figure 7(a).

**Figure 7.**
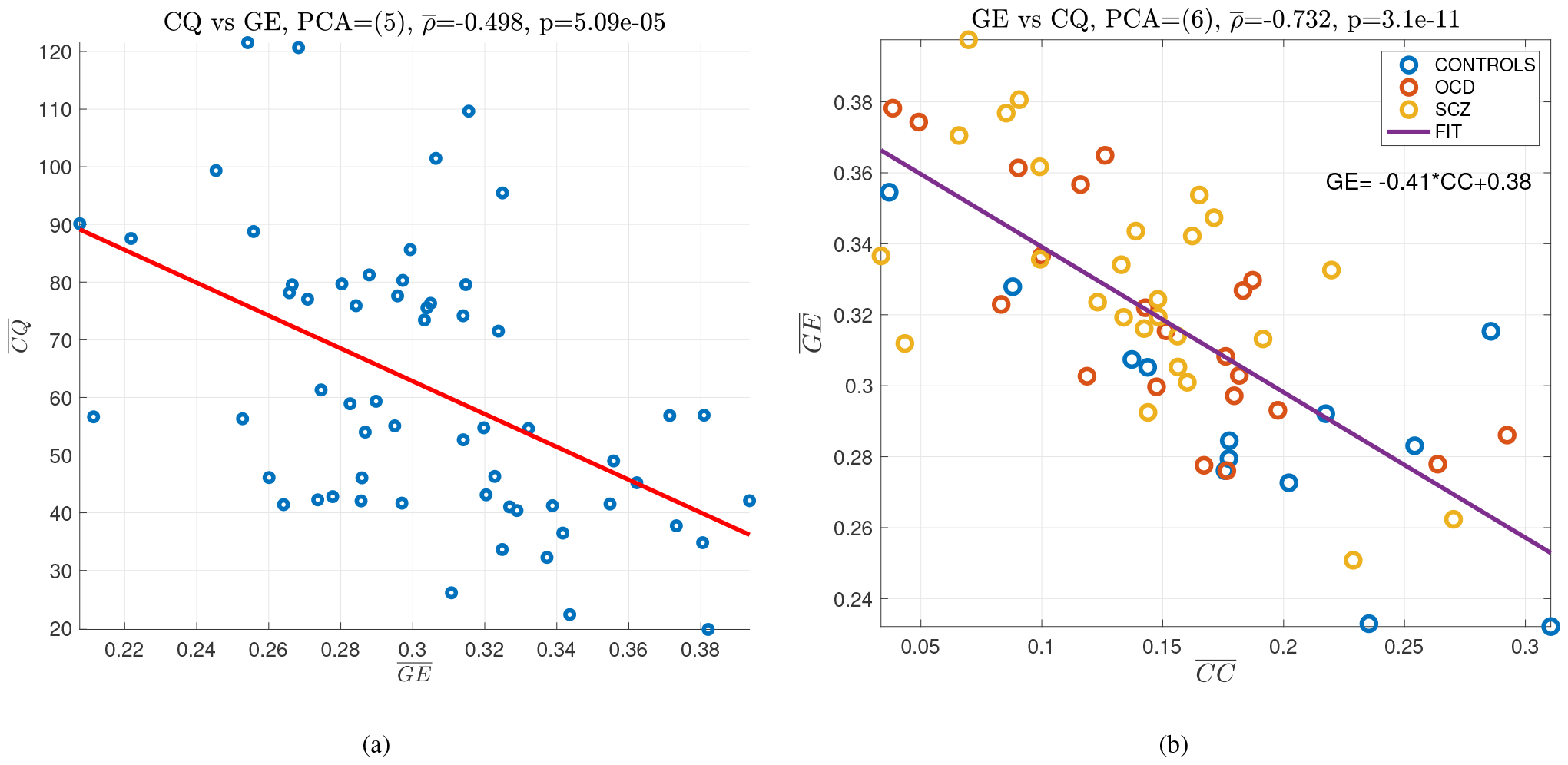
Correlation of a) 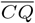 and 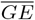 values, and b) 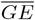 and 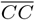 values.

Speaking purely according to neuropsychiatric research, patients with OCD and SCZ exhibit many symptoms in common. Their behavioral results (minds) also support this claim, as evidenced by the similarity of the CQ values shown in Figure 4(b). However, their brains behave differently, as shown in Figures 5. The combination of these mind-body features yields far superior performance than either alone. It appears that it may be better to consider behavioral and physiological data together rather than attempting to find a causal explanation for behavioral findings using correlation analysis.

Figure 7(b)) shows an interesting but predictable result: higher *CC* networks have lower *GE* values. As the *CC* increases in a network, closer nodes are favored to communicate information, resulting in an increase in the number of shorter paths and lowering the global efficiency value.

## DISCUSSIONS

### Behavioral Results

Previous research has yielded conflicting results when investigating various executive tasks in OCD. Some studies have found that OCD subjects make more errors and respond slower in Stroop incongruent trials than control group (Hartston & Swerdlow, 1999; Martinot et al., 1990), while others have found no such differences (Hollander et al., 1993; Schmidtke, Schorb, Winkelmann, & Hohagen, 1998). Patients with SCZ (Lieberman, 1999; Pérez-Neri, Ramírez-Bermúdez, Montes, & Ríos, 2006) and OCD experience cortical neuronal degeneration that correlates with the duration of the disease (Cummings, 2003; Graybiel & Rauch, 2000). In this study, we discovered that individuals with OCD had lower cognitive inhibition than healthy controls. These findings indicate that these impairments may contribute to repetitive symptomatic behaviors associated with OCD, such as compulsion and obsessions.

Neuropsychological assessments revealed disrupted performance associated with frontal lobe functions in both SCZ and OCD, with SCZ showing a more severe deficit. In most cases, SCZ patients suffer more than patients with OCD in terms of executive functions (Tükel et al., 2012; Whitney, Fastenau, Evans, & Lysaker, 2004). This suggests a link between the severity of negative symptoms and SCZ. More specifically, subjects with both disorders made more errors and took longer to complete the Stroop incongruent trials than controls. Simultaneously, lack of blood flow also decreases, resulting in cognitive impairments, as depicted in the 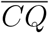 values shown in Figure 4(a).

Our behavioral results shown in Figure 4(b) are consistent with the literature (Chan, Shum, Toulopoulou, & Chen, 2008; Moritz et al., 2002). Specifically, our findings show that patients with SCZ had the poorest performance, while patients with OCD had a less severe but still impaired performance, in contrast to the relatively better performance observed in healthy control subjects.

The current findings show that the Stroop task assessment, which is known for its sensitivity to frontal functioning, found impaired frontal lobe function in both SCZ and OCD, with SCZ showing a more significant impairment. However, one should exercise caution when interpreting this finding because the number of controls (*N* = 13) and affected patients (*N* = 47) is not equal. Given that the variance was expected to be lower in the healthy control group, this may not be considered a significant statistical concern.

### Small-Worldness Results

Small-worldness is a phenomenon discovered in the anatomical and functional connectivity patterns of human brains (Bassett & Bullmore, 2006, 2017). In fact, this has been demonstrated both theoretically and experimentally in several studies (Park, Kim, Kim, & Kim, 2008). Several studies have found a shift toward randomness in the functional connectivity networks of patients with SCZ (Alexander-Bloch et al., 2010; Kambeitz et al., 2016).

Bassett et al. used global efficiency to assess the small-worldness of people with SCZ and healthy controls where they observed that controls. They observed that controls’ *GE* values lie between random and regular networks in the *γ*-band network while SCZ patients have *GE* values closer to regular matrices. However, this pattern reverses in other EEG bands (Bassett et al., 2009). Interestingly, they also discovered a higher *GE* value at the *β*-band for patients with SCZ. Our results also show that psychiatric patients have higher *GE* values than controls (See Figure 5(a)). This is due to the increased cost of operating the network with more but shorter connections. However, there appears to be a methodological conflict with the cost-efficiency analysis. When a matrix is thresholded with the greatest number of connections, the same cost efficiency (density or sparsity) is found regardless of the actual matrix entries. Similarly, in an fMRI study, Yu et al. discovered evidence that schizophrenics have lower *GE* than healthy controls (Q. Yu et al., 2011). This finding contradicts our previous findings and the dissociative nature of the disease. In contrast, Sun et al. discovered that fMRI data increased global efficiency, “indicating a tendency toward a more random organization of temporal brain networks” (Sun, Collinson, Suckling, & Sim, 2019), which is similar to our finding. Their difference from the literature is attributed partly to the choice of data analysis algorithm and a compensatory effort by the schizophrenic brain. In a resting fMRI study, Li et al. found higher *GE* and *σ* values for OCD patients compared to healthy controls (Li, Li, Cao, et al., 2022). Similarly, several studies have shown that *GE* from resting-state fMRI functional connectivity maps is higher than in healthy controls (Sun et al., 2019; M. Yu et al., 2017).

Hence, there is strong evidence that the neuroimaging method chosen, the preprocessing steps, and the algorithm used to compute the small-world features all have an impact on the findings and their interpretation. We propose an algorithm that avoids the pitfalls of previous studies by claiming that 1) there is an inherent background connectivity obscuring the cognition-related topology and it needs to be removed in advance, 2) further cleaning of the connectivity matrices is required to lower the variability; thus, PCA should be preferred, 3) thresholding should be performed by keeping a certain number of the highest values for an individual matrix to ensure the same sparsity for each matrix. Once all of these steps have been completed, we can begin computing the small-world features and determining their statistical significance.

These findings add to the previously known characteristics of people suffering from such diseases. It is well-documented that patients with these diseases exhibit dissociative symptoms in their daily lives. This study sheds light on the impairments in these patients’ functional connectivity patterns, allowing us to determine the cause of these dissociative behavioral symptoms. The increase in GE values compared to healthy controls, as well as the decrease in *CC* and *Q*, all indicate somewhat random (or chaotic) brain patterns. Future studies can use this analysis pipeline to develop biomarkers (or neuromarkers) that can better quantify the severity of functional connectivity pattern impairments.

### Connectivity Patterns

Laterality is a measurement of hemispheric symmetry. We discovered a higher left-sided dominance of psychiatric patients than healthy controls. Similar results were obtained with fNIRS analysis, which revealed a left dominance for the Stroop Task (Aleksandrowicz et al., 2020; Ehlis, Herrmann, Wagener, & Fallgatter, 2005; Koray Ciftci et al., 2008).

### Neuroscientific Results

According to neuropsychiatric research, patients with OCD and SCZ exhibit many symptoms in common. Their behavioral results (minds) support this claim, as evidenced by the similarity of the *CQ* values shown in Figure 4(b). However, their brains behave differently, as shown in Figures 5. The combination of these mind-body features yields far superior performance than either alone. It appears that it may be better to consider behavioral and physiological data together rather than attempting to find a causal explanation for behavioral findings using correlation analysis.

No single neuropathological marker has been identified for either SCZ or OCD. The relationship between the neuroanatomical and neuropsychological mechanisms underlying SCZ and OCD is complex. Although both disorders exhibit frontal lobe pathology, functional neuroimaging studies typically show prefrontal hypoactivity in SCZ (Hazlett et al., 2000) and increased activity in the prefrontal cortex in OCD (Machlin et al., 1991). Hence, it appears logical that in the search for sensitive features, markers focusing on frontal brain regions should be investigated. Furthermore, collecting fNIRS data from the prefrontal cortex is an important and precise method for distinguishing between these disorders. It is essential to acknowledge that although frontal lobe pathology is common to both SCZ and OCD, the underlying neural circuits may involve different structures and neurotransmitter systems. For example, in SCZ, hypofrontality may be associated with a dopaminergic deficit in the frontal cortex (Andreasen et al., 1999), whereas OCD is primarily associated with serotonergic dysregulation in the orbitofrontal cortex (Machlin et al., 1991; Saxena & Rauch, 2000). Given that the Stroop task has been shown to elicit activations in the dorsolateral prefrontal cortex and anterior cingulate (Scarpina & Tagini, 2017; Schroeter, Zysset, Wahl, & von Cramon, 2004; Silton et al., 2010; Zhang, Sun, Sun, Luo, & Gong, 2014; Zysset et al., 2001), it may provide useful insights but might not be sufficient for a comprehensive understanding of the differences in frontal lobe pathology in these disorders.

Figure 7(b) demonstrates a small-world phenomenon, with higher *CC* networks having lower *GE* values. Hence, there appears to be a trade-off between *CC* and *GE* for a network to communicate efficiently, as evidenced by the lower *GE* and higher *CC* values of healthy controls compared to those of people suffering from OCD and SCZ, as shown in Figure 7(b). It might be appropriate to refer to global efficiency as **global cost** because it is a measure of the extensive effort required to communicate information over longer distances.

### Limitations of the study

fNIRS signals contain a wealth of information, of which only a small portion was used in this study. Temporal features of these signals (*e*.*g*. mean, amplitude variations, temporal delays) were not used in the study. These temporal and local values extracted from single channels are known to be closely related to the neurobiology and neurovascular dynamics of the task at hand, as long as these signals are free of contamination from surface fluctuations or other task-unrelated systemic fluctuations. A large body of research has focused on this “cleaning” operation to achieve task-related dynamics. Due to a lack of agreement on the best signal processing pipeline for this problem, this paper concentrated more on overall collective dynamics rather than single channel changes. Therefore, further studies must include these single-channel features to improve the clinical decision-making process, as in (Huang et al., 2018; Ménoret, Farrugia, Pasdeloup, & Gripon, 2017; Ortega, Frossard, Kovačević, Moura, & Vandergheynst, 2018).

To extract meaningful features from a complex dataset, a pipeline of data manipulation steps is required. There are two methods for determining this optimal pipeline: 1) exhaustive search (data-driven) and 2) model-based (hypothesis-driven). In this study, we chose the second approach. This method also necessitates a basic understanding of the model under investigation (i.e. how the brain works). There could be alternatives to the assumptions made in this paper. For example, in our previous paper (Akin, 2021), no background subtraction was performed because the PCA-based approach was assumed to eliminate all background task-unrelated connectivity topologies.

## CONCLUSION

A pipeline for data processing from the fNIRS modality is proposed to demonstrate that people with OCD and SCZ exhibit small-world characteristics similar to random networks. We used a back-subtraction method to eliminate the common network topology, which allowed us to better understand this shift toward randomness. The typical characteristics of small-worldness (*GE, CC, Q*, and *σ*) indicated that the connectivity patterns of these patients were similar to random networks. This means that running the same operation as healthy controls costs more, as evidenced by higher *GE* values and lower *CC* values. To our knowledge, this is the first study to use the fNIRS modality to compare the small-world features of peo

## ACKNOWLEDGMENTS

This project was sponsored by a grant from TUBITAK Project No: 108S101 and Bogazici University Research Fund (BURF Project No: 5106). The authors wish to thank Deniz Nevşehirli for his contributions to fNIRS optode development, Dr. Nermin Topaloĝu and Dr. Sinem Burcu Erdoğan for data collection. We acknowledge the collaboration of Dr. Hasan Herken, Gülfizar Varna in patient recruitment, hypothesis generation, and discussion of the findings.

## REFERENCES

Achard, S., & Bullmore, E. T. (2007). Efficiency and cost of economical brain functional networks. PLoS Comput. Biol., 3(2), 174–183. doi: 10.1371/journal.pcbi.0030017

Adolphs, R. (2003a). Cognitive neuroscience of human social behaviour. Nat Rev Neurosci, 4(3), 165–178. Retrieved from 10.1038/nrn1056 doi: 10.1038/nrn1056

Adolphs, R. (2003b). Investigating the cognitive neuroscience of social behavior. Neuropsychologia, 41(2), 119–126.

Adolphs, R. (2009). The social brain: neural basis of social knowledge. Annu Rev Psychol, 60, 693–716. Retrieved from 10.1146/annurev.psych.60.110707.163514 doi: 10.1146/annurev.psych.60.110707.163514

Akgul, C. B., Akin, A., & Sankur, B. (2006). Extraction of cognitive activity-related waveforms from functional near-infrared spectroscopy signals. Med Biol Eng Comput, 44(11), 945–958. Retrieved from 10.1007/s11517-006-0116-3 doi: 10.1007/s11517-006-0116-3

Akin, A. (2017). Partial correlation-based functional connectivity analysis for functional near-infrared spectroscopy signals. J Biomed Opt, 22(12), 1–10. doi: 10.1117/1.JBO.22.12.126003

Akin, A. (2021). fnirs-derived neurocognitive ratio as a biomarker for neuropsychiatric diseases. Neurophotonics, 8(3). doi: 10.1117/1.NPh.8.3.035008

Aleksandrowicz, A., Hagenmuller, F., Haker, H., Heekeren, K., Theodoridou, A., Walitza, S., … Kawohl, W. (2020). Frontal brain activity in individuals at risk for schizophrenic psychosis and bipolar disorder during the emotional stroop task - an fnirs study. Neuroimage Clin, 26, 102232. doi: 10.1016/j.nicl.2020.102232

Alexander-Bloch, A. F., Gogtay, N., Meunier, D., Birn, R., Clasen, L., Lalonde, F., … Bullmore, E. T. (2010). Disrupted modularity and local connectivity of brain functional networks in childhood-onset schizophrenia. Front Syst Neurosci, 4, 147. doi: 10.3389/fnsys.2010.00147

Andreasen, N. C., Nopoulos, P., O’Leary, D. S., Miller, D. D., Wassink, T., & Flaum, M. (1999). Defining the phenotype of schizophrenia: cognitive dysmetria and its neural mechanisms. Biological psychiatry, 46(7), 908–920.

Ata Akin, Didem Bilensoy, Uzay E Emir, Murat Gulsoy, Selçuk Candansayar, & Hayrunnisa Bolay. (2006). Cerebrovascular dynamics in patients with migraine: near-infrared spectroscopy study. Neurosci Lett, 400(1-2), 86–91. Retrieved from 10.1016/j.neulet.2006.02.016 doi: 10.1016/j.neulet.2006.02.016

Aydöre, S., Mihçak, M. K., Çifti, K., & Akin, A. (2010). On temporal connectivity of PFC via Gauss-Markov modeling of fNIRS signals. Biomedical Engineering, IEEE Transactions on, 57(3), 761–768.

Bassett, D. S., & Bullmore, E. (2006). Small-world brain networks. Neuroscientist, 12(6), 512–23. doi: 10.1177/1073858406293182

Bassett, D. S., & Bullmore, E. T. (2017). Small-world brain networks revisited. Neuroscientist, 23(5), 499–516. doi: 10.1177/1073858416667720

Bassett, D. S., Bullmore, E. T., Meyer-Lindenberg, A., Apud, J. A., Weinberger, D. R., & Coppola, R. (2009). Cognitive fitness of cost-efficient brain functional networks. Proc Natl Acad Sci U S A, 106(28), 11747–52. doi: 10.1073/pnas.0903641106

Bullmore, E., & Sporns, O. (2009). Complex brain networks: graph theoretical analysis of structural and functional systems. Nat Rev Neurosci, 10(3), 186–198. Retrieved from 10.1038/nrn2575 doi: 10.1038/nrn2575

Chan, R. C., Shum, D., Toulopoulou, T., & Chen, E. Y. (2008). Assessment of executive functions: Review of instruments and identification of critical issues. Archives of clinical neuropsychology, 23(2), 201–216.

Ciftçi, K., Sankur, B., Kahya, Y. P., & Akin, A. (2008). Multilevel statistical inference from functional near-infrared spectroscopy data during stroop interference. IEEE Trans Biomed Eng, 55(9), 2212–20. doi: 10.1109/TBME.2008.923918

Cummings, J. L. (2003). Toward a molecular neuropsychiatry of neurodegenerative diseases. Annals of neurology, 54(2), 147–154.

Dadgostar, M., Setarehdan, S. K., Shahzadi, S., & Akin, A. (2016). Functional connectivity of the pfc via partial correlation. Optik-International Journal for Light and Electron Optics, 127(11), 4748–4754.

Dalmis, M. U., & Akin, A. (2015). Similarity analysis of functional connectivity with functional near-infrared spectroscopy. J Biomed Opt, 20(8), 86012. doi: 10.1117/1.JBO.20.8.086012

Deuker, L., Bullmore, E. T., Smith, M., Christensen, S., Nathan, P. J., Rockstroh, B., & Bassett, D. S. (2009). Reproducibility of graph metrics of human brain functional networks. Neuroimage, 47(4), 1460–1468. Retrieved from 10.1016/j.neuroimage.2009.05.035 doi: 10.1016/j.neuroimage.2009.05.035

Ehlis, A.-C., Herrmann, M. J., Wagener, A., & Fallgatter, A. J. (2005). Multi-channel near-infrared spectroscopy detects specific inferior-frontal activation during incongruent Stroop trials. Biol Psychol, 69(3), 315–331. Retrieved from 10.1016/j.biopsycho.2004.09.003 doi: 10.1016/j.biopsycho.2004.09.003

Einalou, Z., Maghooli, K., Setarehdan, S. K., & Akin, A. (2014). Functional near infrared spectroscopy to investigation of functional connectivity in schizophrenia using partial correlation. Universal Journal of Biomedical Engineering, 2(1), 5–8.

Einalou, Z., Maghooli, K., Setarehdan, S. K., & Akin, A. (2015). Functional near Infrared Spectroscopy for Functional Connectivity during Stroop test via Mutual Information. Adv. Biores., 6(1), 62–67.

Einalou, Z., Maghooli, K., Setarehdan, S. K., & Akin, A. (2016). Effective channels in classification and functional connectivity pattern of prefrontal cortex by functional near infrared spectroscopy signals. Optik-International Journal for Light and Electron Optics, 127(6), 3271–3275.

Einalou, Z., Maghooli, K., Setarehdan, S. K., & Akin, A. (2017). Graph theoretical approach to functional connectivity in prefrontal cortex via fnirs. Neurophotonics, 4(4), 041407. doi: 10.1117/1.NPh.4.4.041407

Elgammal, A., Harwood, D., & Davis, L. (2000). Non-parametric model for background subtraction. In European conference on computer vision (pp. 751–767).

Erdoĝn, S. B., Yücel, M. A., & Akın, A. (2014). Analysis of task-evoked systemic interference in fNIRS measurements: insights from fMRI. Neuroimage, 87, 490–504. Retrieved from 10.1016/j.neuroimage.2013.10.024 doi: 10.1016/j.neuroimage.2013.10.024

Erdoğan, S. B., & Yükselen, G. (2022). Four-class classification of neuropsychiatric disorders by use of functional near-infrared spectroscopy derived biomarkers. Sensors, 22(14), 5407.

Fox, M. D., Snyder, A. Z., Vincent, J. L., Corbetta, M., Van Essen, D. C., & Raichle, M. E. (2005). The human brain is intrinsically organized into dynamic, anticorrelated functional networks. Proc. Natl. Acad. Sci. U. S. A., 102(27), 9673–9678. doi: 10.1073/pnas.0504136102

Gao, Z., Xiao, Y., Zhu, F., Tao, B., Yu, W., & Lui, S. (2023). The whole-brain connectome landscape in patients with schizophrenia: A systematic review and meta-analysis of graph theoretical characteristics. Neurosci Biobehav Rev, 148, 105144. doi: 10.1016/j.neubiorev.2023.105144

Graybiel, A. M., & Rauch, S. L. (2000). Toward a neurobiology of obsessive-compulsive disorder. Neuron, 28(2), 343–347.

Hartston, H. J., & Swerdlow, N. R. (1999). Visuospatial priming and stroop performance in patients with obsessive compulsive disorder. Neuropsychology, 13(3), 447–57. doi: 10.1037//0894-4105.13.3.447

Hazlett, E. A., Buchsbaum, M. S., Jeu, L. A., Nenadic, I., Fleischman, M. B., Shihabuddin, L., … Harvey, P. D. (2000). Hypofrontality in unmedicated schizophrenia patients studied with pet during performance of a serial verbal learning task. Schizophr Res, 43(1), 33–46. doi: 10.1016/s0920-9964(99)00178-4

He, Y., & Evans, A. (2010). Graph theoretical modeling of brain connectivity. Curr Opin Neurol, 23(4), 341–350. Retrieved from 10.1097/WCO.0b013e32833aa567 doi: 10.1097/WCO.0b013e32833aa567

Hofman, M. A. (2012). Design principles of the human brain: an evolutionary perspective. Prog Brain Res, 195, 373–90. doi: 10.1016/B978-0-444-53860-4.00018-0

Hofman, M. A. (2014). Evolution of the human brain: when bigger is better. Front Neuroanat, 8, 15. doi: 10.3389/fnana.2014.00015

Hollander, E., Cohen, L., Richards, M., Mullen, L., DeCaria, C., & Stern, Y. (1993). A pilot study of the neuropsychology of obsessive-compulsive disorder and parkinson’s disease: basal ganglia disorders. J Neuropsychiatry Clin Neurosci, 5(1), 104–7. doi: 10.1176/jnp.5.1.104

Horprasert, T., Harwood, D., & Davis, L. S. (1999). A statistical approach for real-time robust background subtraction and shadow detection. In Ieee iccv (Vol. 99, pp. 1–19).

Huang, W., Bolton, T. A., Medaglia, J. D., Bassett, D. S., Ribeiro, A., & Van De Ville, D. (2018). A graph signal processing perspective on functional brain imaging. Proceedings of the IEEE, 106(5), 868–885.

Kambeitz, J., Kambeitz-Ilankovic, L., Cabral, C., Dwyer, D. B., Calhoun, V. D., van den Heuvel, M. P., … Malchow, B. (2016). Aberrant functional whole-brain network architecture in patients with schizophrenia: A meta-analysis. Schizophr Bull, 42 Suppl 1(Suppl 1), S13–21. doi: 10.1093/schbul/sbv174

Klados, M. A., Kanatsouli, K., Antoniou, I., Babiloni, F., Tsirka, V., Bamidis, P. D., & Micheloyannis, S. (2013). A graph theoretical approach to study the organization of the cortical networks during different mathematical tasks. PLoS One, 8(8), e71800. Retrieved from 10.1371/journal.pone.0071800 doi: 10.1371/journal.pone.0071800

Koray Ciftci, Bulent Sankur, Yasemin P Kahya, & Ata Akin. (2008). Multilevel statistical inference from functional near-infrared spectroscopy data during stroop interference. IEEE Trans Biomed Eng, 55(9), 2212–2220. Retrieved from 10.1109/TBME.2008.923918 doi: 10.1109/TBME.2008.923918

Li, X., Li, H., Cao, L., Liu, J., Xing, H., Huang, X., & Gong, Q. (2022). Application of graph theory across multiple frequency bands in drug-näive obsessive-compulsive disorder with no comorbidity. J Psychiatr Res, 150, 272–278. doi: 10.1016/j.jpsychires.2022.03.041

Li, X., Li, H., Jiang, X., Li, J., Cao, L., Liu, J., … Gong, Q. (2022). Characterizing multiscale modular structures in medication-free obsessive-compulsive disorder patients with no comorbidity. Hum Brain Mapp, 43(7), 2391–2399. doi: 10.1002/hbm.25794

Lieberman, J. A. (1999). Is schizophrenia a neurodegenerative disorder? a clinical and neurobiological perspective. Biol Psychiatry, 46(6), 729–39. doi: 10.1016/s0006-3223(99)00147-x

Machlin, S. R., Harris, G. J., Pearlson, G. D., Hoehn-Saric, R., Jeffery, P., & Camargo, E. E. (1991). Elevated medial-frontal cerebral blood flow in obsessive-compulsive patients: a spect study. Am J Psychiatry, 148(9), 1240–2. doi: 10.1176/ajp.148.9.1240

Martinot, J., Allilaire, J., Mazoyer, B., Hantouche, E., Huret, J., Legaut-Demare, F., … others (1990). Obsessive-compulsive disorder: A clinical, neuropsychological and positron emission tomography study. Acta Psychiatrica Scandinavica, 82(3), 233–242.

Mènoret, M., Farrugia, N., Pasdeloup, B., & Gripon, V. (2017). Evaluating graph signal processing for neuroimaging through classification and dimensionality reduction. In 2017 ieee global conference on signal and information processing (globalsip) (pp. 618–622).

Moritz, S., Birkner, C., Kloss, M., Jahn, H., Hand, I., Haasen, C., & Krausz, M. (2002). Executive functioning in obsessive–compulsive disorder, unipolar depression, and schizophrenia. Archives of Clinical Neuropsychology, 17(5), 477–483.

Nandi, D., Ashour, A. S., Samanta, S., Chakraborty, S., Salem, M. A., & Dey, N. (2015). Principal component analysis in medical image processing: a study. International Journal of Image Mining, 1(1), 65–86.

Niu, H., Li, Z., Liao, X., Wang, J., Zhao, T., Shu, N., … He, Y. (2013). Test-retest reliability of graph metrics in functional brain networks: a resting-state fnirs study. PLoS One, 8(9), e72425. doi: 10.1371/journal.pone.0072425

Oliver, N. M., Rosario, B., & Pentland, A. P. (2000). A bayesian computer vision system for modeling human interactions. IEEE transactions on pattern analysis and machine intelligence, 22(8), 831–843.

Ortega, A., Frossard, P., Kovac?ević, J., Moura, J. M., & Vandergheynst, P. (2018). Graph signal processing: Overview, challenges, and applications. Proceedings of the IEEE, 106(5), 808–828.

Palva, S., Monto, S., & Palva, J. M. (2010). Graph properties of synchronized cortical networks during visual working memory maintenance. Neuroimage, 49(4), 3257–3268. Retrieved from 10.1016/j.neuroimage.2009.11.031 doi: 10.1016/j.neuroimage.2009.11.031

Park, C.-h., Kim, S. Y., Kim, Y.-H., & Kim, K. (2008). Comparison of the small-world topology between anatomical and functional connectivity in the human brain. Physica A: statistical mechanics and its applications, 387(23), 5958–5962.

Pèrez-Neri, I., Ramírez-Bermúdez, J., Montes, S., & Ríos, C. (2006). Possible mechanisms of neurodegeneration in schizophrenia. Neurochem Res, 31(10), 1279–94. doi: 10.1007/s11064-006-9162-3

Piccardi, M. (2004). Background subtraction techniques: a review. In 2004 ieee international conference on systems, man and cybernetics (ieee cat. no. 04ch37583) (xVol. 4, pp. 3099–3104).

Raichle, M. E., MacLeod, A. M., Snyder, A. Z., Powers, W. J., Gusnard, D. A., & Shulman, G. L. (2001). A default mode of brain function. Proc. Natl. Acad. Sci. U. S. A., 98(2), 676–682.

Ralchle, M. E., & Snyder, A. Z. (2007). A default mode of brain function: A brief history of an evolving idea. Neuroimage, 37(4), 1083–1090. doi: 10.1016/j.neuroimage.2007.02.041

Rodarmel, C., & Shan, J. (2002). Principal component analysis for hyperspectral image classification. Surveying and Land Information Science, 62(2), 115–122.

Rubinov, M., & Sporns, O. (2010). Complex network measures of brain connectivity: uses and interpretations. Neuroimage, 52(3), 1059–1069.

Saxena, S., & Rauch, S. L. (2000). Functional neuroimaging and the neuroanatomy of obsessive-compulsive disorder. Psychiatr Clin North Am, 23(3), 563–86. doi: 10.1016/s0193-953x(05)70181-7

Scarpina, F., & Tagini, S. (2017). The stroop color and word test. Frontiers in psychology, 8, 557.

Schmidtke, K., Schorb, A., Winkelmann, G., & Hohagen, F. (1998). Cognitive frontal lobe dysfunction in obsessive-compulsive disorder. Biol Psychiatry, 43(9), 666–73. doi: 10.1016/s0006-3223(97)00355-7

Schroeter, M. L., Zysset, S., Wahl, M., & von Cramon, D. Y. (2004). Prefrontal activation due to Stroop interference increases during development–an event-related fNIRS study. Neuroimage, 23(4), 1317–1325. Retrieved from 10.1016/j.neuroimage.2004.08.001 doi: 10.1016/j.neuroimage.2004.08.001

Sergul Aydore, M. Kivanc Mihcak, Koray Ciftci, & Ata Akin. (2010). On temporal connectivity of pfc via gauss-markov modeling of fnirs signals. IEEE Trans Biomed Eng, 57(3), 761–768. Retrieved from 10.1109/TBME.2009.2020792 doi: 10.1109/TBME.2009.2020792

Silton, R. L., Heller, W., Towers, D. N., Engels, A. S., Spielberg, J. M., Edgar, J. C., … Miller, G. A. (2010). The time course of activity in dorsolateral prefrontal cortex and anterior cingulate cortex during top-down attentional control. NeuroImage, 50, 1292–1302. doi: 10.1016/j.neuroimage.2009.12.061

Skidmore, F., Korenkevych, D., Liu, Y., He, G., Bullmore, E., & Pardalos, P. M. (2011). Connectivity brain networks based on wavelet correlation analysis in parkinson fMRI data. Neurosci Lett, 499(1), 47–51. Retrieved from 10.1016/j.neulet.2011.05.030 doi: 10.1016/j.neulet.2011.05.030

Sun, Y., Collinson, S. L., Suckling, J., & Sim, K. (2019). Dynamic reorganization of functional connectivity reveals abnormal temporal efficiency in schizophrenia. Schizophr Bull, 45(3), 659–669. doi: 10.1093/schbul/sby077

Thorsen, A. L., Vriend, C., de Wit, S. J., Ousdal, O. T., Hagen, K., Hansen, B., … van den Heuvel, O. A. (2021). Effects of bergen 4-day treatment on resting-state graph features in obsessive-compulsive disorder. Biol Psychiatry Cogn Neurosci Neuroimaging, 6(10), 973–982. doi: 10.1016/j.bpsc.2020.01.007

Tükel, R., Gürvit, H., Ertekin, B. A., Oflaz, S., Ertekin, E., Baran, B., … Atalay, F. (2012). Neuropsychological function in obsessive-compulsive disorder. Comprehensive psychiatry, 53(2), 167–175.

Whitney, K. A., Fastenau, P. S., Evans, J. D., & Lysaker, P. H. (2004). Comparative neuropsychological function in obsessive-compulsive disorder and schizophrenia with and without obsessive-compulsive symptoms. Schizophrenia research, 69(1), 75–83.

Xia, Y., Wen, L., Eberl, S., Fulham, M., & Feng, D. (2008). Genetic algorithm-based pca eigenvector selection and weighting for automated identification of dementia using fdg-pet imaging. Annu Int Conf IEEE Eng Med Biol Soc, 2008, 4812–5. doi: 10.1109/IEMBS.2008.4650290

Yeo, B. T. T., Krienen, F. M., Sepulcre, J., Sabuncu, M. R., Lashkari, D., Hollinshead, M., … Buckner, R. L. (2011). The organization of the human cerebral cortex estimated by intrinsic functional connectivity. J Neurophysiol, 106(3), 1125–65. doi: 10.1152/jn.00338.2011

Yu, M., Dai, Z., Tang, X., Wang, X., Zhang, X., Sha, W., … Zhang, Z. (2017). Convergence and divergence of brain network dysfunction in deficit and non-deficit schizophrenia. Schizophr Bull, 43(6), 1315–1328. doi: 10.1093/schbul/sbx014

Yu, Q., Sui, J., Rachakonda, S., He, H., Pearlson, G., & Calhoun, V. D. (2011). Altered small-world brain networks in temporal lobe in patients with schizophrenia performing an auditory oddball task. Front Syst Neurosci, 5, 7. doi: 10.3389/fnsys.2011.00007

Zhang, L., Sun, J., Sun, B., Luo, Q., & Gong, H. (2014). Studying hemispheric lateralization during a stroop task through near-infrared spectroscopy-based connectivity. J Biomed Opt, 19(5), 57012. Retrieved from 10.1117/1.JBO.19.5.057012 doi: 10.1117/1.JBO.19.5.057012

Zysset, S., Müller, K., Lohmann, G., & von Cramon, D. Y. (2001). Color-word matching Stroop task: separating interference and response conflict. Neuroimage, 13(1), 29–36. Retrieved from 10.1006/nimg.2000.0665 doi: 10.1006/nimg.2000.0665

